# *In vitro* evaluation of protein–protein interactions in the rice KAI2 ligand signaling complex

**DOI:** 10.1101/2025.09.29.679124

**Authors:** Keita Tanaka, Jiawang Wu, Qianwei Xia, Yutaro Harada, Taiki Suzuki, Yujiao Yan, Yoshiya Seto, Guosheng Xiong, Hiromu Kameoka

## Abstract

KARRIKIN INSENSITIVE 2 (KAI2)/DWARF14-LIKE (D14L) plays key roles in land plant development, environmental responses, and the establishment of arbuscular mycorrhizal symbiosis, likely acting as the receptor for unidentified signaling molecules termed KAI2 ligands (KLs). KL perception by KAI2/D14L promotes DWARF3 (D3)/MORE AXILLARY GROWTH2 (MAX2) F-box protein-mediated ubiquitination of SUPPRESSOR OF MAX2 1 (SMAX1) proteins, thereby transducing the KL signals. Although genetic and *in vivo* assays have demonstrated the functions of these components, the biochemical details of their interactions remain elusive. Here we investigated physical interactions between rice D14L, D3, and OsSMAX1 *in vitro* using desmethyl germinone (dMGer), a recently developed KL analog. dMGer elicited KL responses in rice with higher activity and pathway specificity than a widely used KL analog (−)-GR24. dMGer, but not (−)-GR24, directly bound to D14L and promoted the interaction between D14L and D3 *in vitro*. The interaction between D14L and OsSMAX1 was also enhanced by dMGer. Furthermore, we identified the domain of OsSMAX1 that distinguishes it from its paralog DWARF53 (D53), which is associated with the strigolactone signaling complex. These findings propose a model of the interactions among KL signaling components and highlight the role of the ligand in the signaling complex.

## Introduction

The α/β-fold hydrolase KARRIKIN INSENSITIVE2 (KAI2)/DWARF14-Like (D14L) is conserved in land plants and regulates diverse aspects of their life cycles. *KAI2* was first identified in *Arabidopsis thaliana* (Arabidopsis) as a gene essential for the response to karrikins (KARs), a group of smoke-derived butenolides that stimulate seed germination and photomorphogenesis (Chiwocha et al., 2009; Waters et al., 2012). *KAI2*/*D14L* regulates not only seed germination and photomorphogenesis but also root development (Villaécija-Aguilar et al., 2019; Carbonnel et al., 2020; Meng et al., 2022), drought response (Li et al., 2017), arbuscular mycorrhizal (AM) symbiosis (Gutjahr et al., 2015), etc. Notably, KARs are absent in living plants, whereas *kai2*/*d14l* mutants exhibit phenotypes opposite to those induced by KAR treatment in wild-type (WT) plants. For example, KAR treatment suppresses mesocotyl elongation in *Oryza sativa* (rice) in the dark, whereas *d14l* mutants exhibit elongated mesocotyls (Gutjahr et al., 2015; Zheng et al., 2020). These findings suggest that KAI2/D14L mediates responses to an unidentified endogenous compound distinct from KARs. Since KAI2/D14L is considered to be its receptor, this putative compound is referred to as KAI2 ligand (KL) (Conn and Nelson, 2016).

The KL signaling pathway is composed of components of the strigolactone (SL) signaling pathway or their paralogs. In SL signaling, SLs bind to DWARF14 (D14)/DECREASED APICAL DOMINANCE2/RAMOSUS3, a paralog of KAI2/D14L, promoting the interaction of D14 with the F-box protein MORE AXILLARY GROWTH2 (MAX2)/DWARF3 (D3) in an Skp, Cullin, F-box (SCF) complex. D14 also interacts with the repressor of the SL signal, DWARF53 (D53) in rice or SUPPRESSOR OF MAX2 1 (SMAX1)-LIKE6/7/8 (SMXL6/7/8) in Arabidopsis, in an SL-dependent manner (Jiang et al., 2013; Zhou et al., 2013; Wang et al., 2015; Wang et al., 2020a; Soundappan et al., 2015). SLs thus trigger the formation of a D14-SCF^MAX2/D3^-D53/SMXL6/7/8 complex and subsequent ubiquitination and 26S-proteasomal degradation of the repressor D53/SMXL6/7/8 proteins, thereby inducing SL responses. On the other hand, in KL signaling, KL is considered to be perceived by KAI2/D14L, which triggers the formation of the KAI2/D14L–SCF^MAX2/D3^–SMAX1/SMXL2/OsSMAX1 complex and their degradation. However, the protein–protein interactions among the KL signaling components have been less well characterized than those in the SL signaling components.

Evidence for the ligand-dependent interactions among the KL signaling components remains limited. In Arabidopsis, although an *in vitro* pull-down assay shows that KAI2 and MAX2 slightly interact in a ligand-dependent manner, this interaction is much weaker than that that observed between AtD14 and MAX2 detected under the same conditions (Xu et al., 2018). Ligand-dependent interaction between KAI2 and SMAX1 has been demonstrated only by *in vivo* co-immunoprecipitation or yeast two-hybrid assays, but not by *in vitro* pull-down assays (Khosla et al., 2020; Yao et al., 2021; Okabe et al., 2023). In rice, ligand-dependent interactions among the KL signaling components have not been reported. Overall, the available evidence is insufficient to determine whether the ligands induce direct interactions among the KL signaling components.

The functions of the domains of the KL signaling components are also poorly understood. SMXL proteins contain conserved domains, N-terminal domain (N), putative ATPase domain 1 (D1), middle region (M), and C-terminal putative ATPase domain 2 (D2). The D2 domain is required for MAX2/D3-mediated degradation (Jiang et al., 2013; Zhou et al., 2013; Wang et al., 2020b; Khosla et al., 2020). Yeast two-hybrid assays have shown that the D1, or D1 and M (D1M) domains are responsible for the ligand-dependent interaction with D14 or KAI2/D14L (Zhou et al., 2013; Khosla et al., 2020). In contrast, another study reported that the D53-D2 domain alone interacts with D14 and that this interaction is enhanced by the C-terminal α-helix of D3 (D3-CTH). Binding of D3-CTH suppresses the ligand-hydrolysis activity of D14 (Shabek et al., 2018). Expression of *MAX2* lacking CTH failed to fully rescue the *max2* mutant phenotypes associated with both SL and KL. Furthermore, mutations in the CTH of D3 or MAX2 showed altered their activity to regulate the hydrolase activity of KAI2 and the SMAX1 ubiquitination, suggesting the involvement of D3/MAX2-CTH in KL signaling (Tal et al., 2022; Tal et al., 2023). However, the role of the CTH in the interactions among the KL signaling components remains to be elucidated.

A possible reason for the limited evidence of the *in vitro* interaction is insufficient activation of KAI2/D14L by KARs or artificial KL analogs. Differential scanning fluorimetry (DSF) has been used to evaluate the activity of the putative ligands of D14 or KAI2/D14L, in which the shifts in thermal stability (melting temperature; Tm) of the proteins indicate structural changes induced by ligand binding (Niesen et al., 2007). Previous works suggest that not ligand binding alone but the shift in receptor stability in DSF is the better indicator of ligand activity (Seto et al., 2019; Kushihara et al., 2025). Natural SLs and (+)-GR24, an artificial SL analog, induce large shifts towards destabilization of D14 (Hamiaux et al., 2012; Abe et al., 2014; de Saint Germain et al., 2016; Seto et al., 2019; Yao et al., 2021) (Supplementary Figure S1). On the other hand, KARs do not induce shifts in KAI2/D14L (Waters et al., 2015; Fukui et al., 2019). This result has raised a hypothesis that the direct ligands of KAI2/D14L are not KARs themselves, but uncharacterized metabolite(s) derived from KARs in plants. Interestingly, (−)-GR24 (Supplementary Figure S1), a widely used artificial KL analog, induces only slight shifts in KAI2 stability in Arabidopsis (Fukui et al., 2019; Yao et al., 2021). Moreover, it does not induce shifts in some species, including rice, *Brachypodium distachyon*, and *Selaginella moellendorffii* (Waters et al., 2015; Yao et al., 2021; Meng et al., 2022). These results suggest the possibility that (−)-GR24 does not possess sufficient activity to enhance the interactions between the KL signaling components.

Recently, a new type of compounds, desmethyl butenolides, has been reported to function as KL analogs (Yao et al., 2021). While natural SLs, artificial SL analogs, and (−)-GR24 have a methyl group on their butenolide ring, desmethyl-type KL analogs lack this methyl group. Currently, four types of desmethyl-type KL analogs, desmethyl Yoshimulactone Green (dYLG, Supplementary Figure S1), desmethyl-GR24 (dGR24), desmethyl-CN-debranone, and desmethyl germinone (dMGer, Supplementary Figure S1), have been developed (Yao et al., 2018; Yao et al., 2021; Okabe et al., 2023). dYLG binds to the ligand binding pocket of KAI2/D14L and emits fluorescence upon hydrolysis by KAI2/D14L. The other three compounds have been shown to induce stronger KL responses *in vivo* and larger stability shifts of KAI2/D14L in DSF than (−)-GR24 (Yao et al., 2021; Meng et al., 2022; Okabe et al., 2023; Kushihara et al., 2025). dMGer reportedly induces the interaction between Arabidopsis KAI2 and SMAX1 in yeast two-hybrid assays (Okabe et al., 2023; Kushihara et al., 2025). However, the ability of desmethyl butenolide-type ligands to induce the *in vitro* interactions of the KL signaling components has not yet been examined.

Here, we investigate the rice KL signaling using artificial KL analogs, (−)-GR24 and dMGer to address key questions: whether these KL analogs directly bind to the receptor and elicit physiological responses across species; whether the KL signaling components directly interact and, if so, which interactions are mediated by KL; and which domain is pivotal for the interactions among the components. Pharmacological assays reveal that dMGer activates the KL signaling pathway more strongly and specifically than (−)-GR24 in rice. *In vitro* assays demonstrate that D14L recognizes dMGer, but not (−)-GR24. We then verified interactions among D14L, D3, and OsSMAX1, providing direct evidence for the enhancement of D14L–D3 and D14L–OsSMAX1 interactions *in vitro*. Furthermore, by assaying the interactions among D14L, D14, and domains of OsSMAX1 and D53, we reveal the functions of the domains that underlie the specific interactions of D14L–OsSMAX1 and D14–D53. Thus, our study refines the biochemical understanding of the KL signaling pathway.

## Results

### dMGer induces KL responses in rice

While the activities of desmethyl-type KL analogs have been assessed with D14L/KAI2 from several species *in vitro*, their KAI2/D14L-dependent activities *in vivo* have been examined only in Arabidopsis. dMGer, a recently developed desmethyl-type KL analog, promotes seed germination and inhibits hypocotyl elongation in Arabidopsis in a KAI2-dependent manner (Okabe et al., 2023; Kushihara et al., 2025). To evaluate the activities of dMGer as a KL analog, we examined the responses of rice to dMGer.

We first assessed the dMGer activity on mesocotyl length. Elongation of mesocotyls in dark-grown rice seedlings is a well-characterized phenotype regulated by the KL signaling pathway (Kameoka and Kyozuka, 2015; Gutjahr et al., 2015; Choi et al., 2020; Zheng et al., 2020). *d14l* mutants exhibit longer mesocotyls than WT, whereas KARs and (−)-GR24 suppress mesocotyl elongation in a *D14L*-dependent manner (Zheng et al., 2020). We compared mesocotyl length of seedlings grown in the dark with or without 1 µM dMGer treatment. WT seedlings treated with dMGer developed shorter mesocotyls than those of the control group (Figure 1A and 1B). In contrast to WT seedlings, *d14l* seedlings showed no significant difference in mesocotyl length between dMGer-treated and untreated groups (Figure 1A and 1B). These results indicate that dMGer suppresses mesocotyl elongation through the KL signaling pathway in rice.

**Figure 1.**
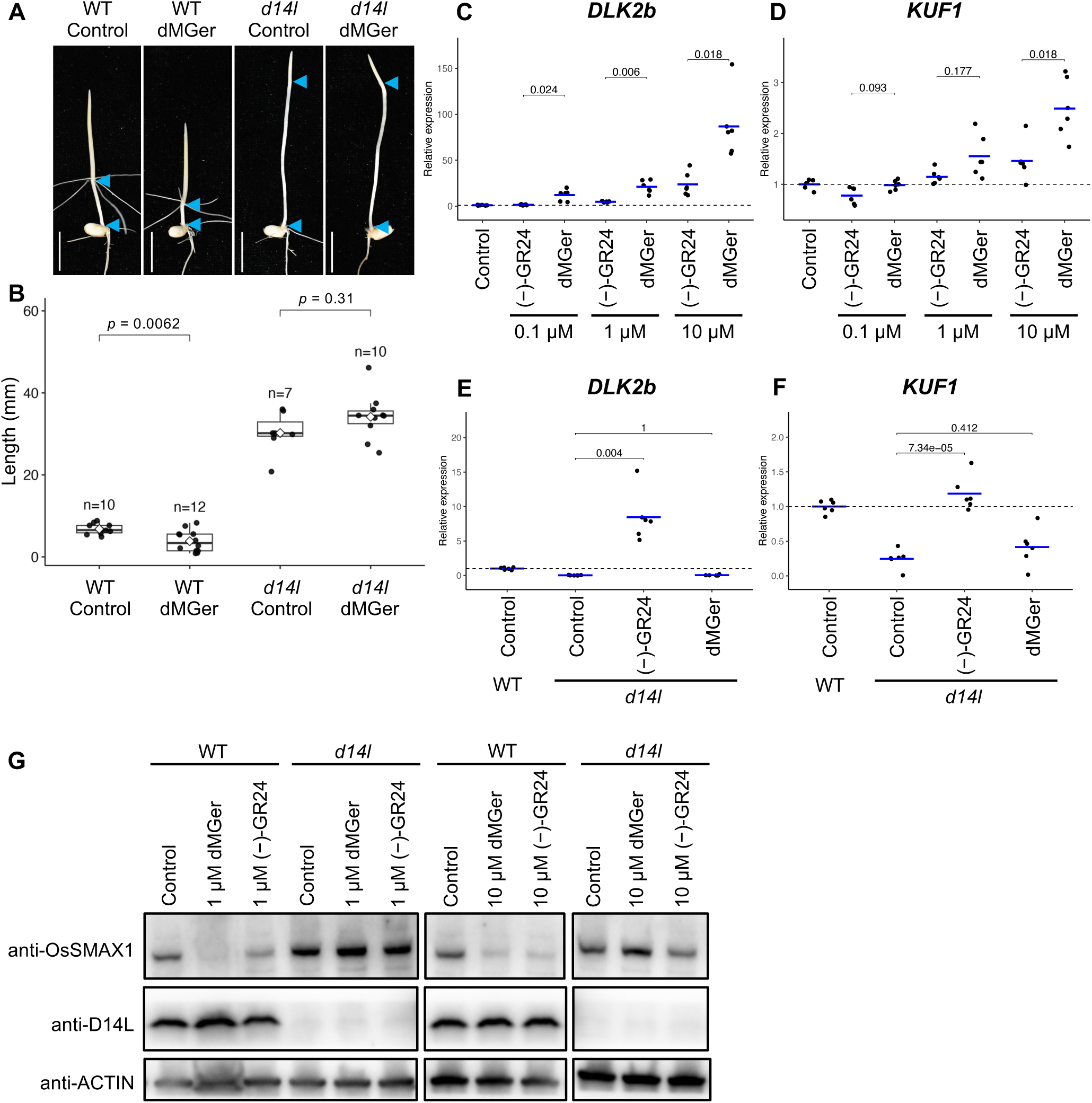
Response of rice seedlings to (−)-GR24 and desmethyl germinone treatments. (**A**) Representative images of 8-day-old wild-type (WT) and *d14l-c2* seedlings treated with 1 µM desmethyl germinone (dMGer) or 0.1% (v/v) acetone as a control. Scale bars: 1 cm. (**B**) Length of mesocotyls of WT and *d14l-c2* plants treated with or without dMGer. Circles represent individual data points. Open squares indicate means of each treatment group. Sample numbers are presented above the plot for each condition. For statistical analysis, Welch’s t-test with Bonferroni correction was performed comparing the control and dMGer-treated groups of each genotype. Calculated *p*-values are presented above the plots for each genotype. (**C**, **D**) Transcript levels of *DLK2b* (**C**) and *KUF1* (**D**) in 5-day-old WT rice roots treated with varying concentrations of (−)-GR24 or dMGer, or with 0.1% (v/v) acetone as control (*n* = 6). Values are expressed as fold change relative to the mean value of the control. Circles represent individual data points; bars indicate means of replicates. For statistical analysis, Welch’s t-test with Bonferroni correction was performed comparing the groups treated with the same concentration of (−)-GR24 or dMGer. Calculated *p*-values are presented above the plots for each pair of the compared groups. (**E**, **F**) Transcript levels of *DLK2b* (**E**) and *KUF1* (**F**) in 5-day-old WT and *d14l-c1* mutant roots treated with 10 µM (−)-GR24 or dMGer (*n* = 6). Acetone (0.1% (v/v)) was used as the control treatment. Values are expressed relative to the mean value of the control WT plants. Circles represent individual data points; bars indicate means of triplicates. For statistical analysis, Welch’s t-test with Bonferroni correction was performed comparing the groups treated with the same concentration of (−)-GR24 or dMGer. Calculated *p*-values are presented above the plots for each pair of the compared groups. (**G**) Levels of OsSMAX1 in calli of WT and *d14l* at two hours following treatment with dMGer or (−)-GR24 at the indicated concentrations. OsSMAX1, D14L, and ACTIN proteins were detected by immunoblotting with anti-OsSMAX1, anti-D14L, and anti-ACTIN antibodies, respectively. Immunobloting with the same antibody was performed on a single membrane. Alt text: Panels labeled A to G showing effects of KAI2 ligand analogs. A presents pictures of rice mesocotyls treated with desmethyl germinone. B is a plot of mesocotyl length, with statistic values. C and D are plots of marker gene expression levels in wild-type rice treated with different concentrations of desmethyl germinone and (−)-GR24, with statistic values. E and F are plots of marker gene expression levels in wild-type and *d14l* mutant rice treated with these KAI2 ligand analogs, with statistic values. G is immunoblot showing effects of the KAI2 ligand analogs on SMAX1 protein levels.

Next, we characterized the effects of dMGer on transcriptional responses of rice KL-responsive markers, *D14-LIKE2b* (*DLK2b*) (Os01g0595600) and *KARRIKIN-UP-REGULATED F-BOX1* (*KUF1*) (Os06g0711700), to dMGer treatment, comparing them with those induced by (−)-GR24. Previous studies have shown that, across land plant species, homologues of these genes are induced by the KL signals (Waters et al., 2012; Choi et al., 2020; Zheng et al., 2020; Mizuno et al., 2021; Sepulveda et al., 2022; Meng et al.). In WT seedlings, dMGer induced the expression of *DLK2b* (Figure 1C) and *KUF1* (Figure 1D) in a dose-dependent manner. Across all concentration tested, dMGer induced transcription of *DLK2b* to a greater extent than (−)-GR24 (Figure 1C). A similar trend was observed for *KUF1* in plants treated with 1 µM and 10 µM of these chemicals (Figure 1D). These results indicate that dMGer induces stronger transcriptional activation of KL-responsive markers in rice than (−)-GR24. To determine whether these responses require *D14L*, we examined the responses in *d14l* mutant seedlings. In *d14l*, 10 µM dMGer did not induce either *DLK2b* (Figure 1E) or *KUF1* (Figure 1F). In contrast, 10 µM (−)-GR24 strongly induced *DLK2b* (Figure 1E). The expression of *KUF1* also tended to be induced by (−)-GR24 (Figure 1F). These results indicate that dMGer induces the expression of these genes depending on *D14L*, whereas (−)-GR24 also acts *D14L*-independently.

Furthermore, we examined the effect of dMGer on OsSMAX1 protein stability *in vivo*. Treatment with 1 µM dMGer led to a marked reduction in OsSMAX1 levels in WT plants within two hours. Consistent with the transcriptional analyses of the KL-responsive marker genes, 1 µM (−)-GR24 also triggered a reduction in OsSMAX1 levels, albeit to a lesser extent than dMGer. In *d14l* mutants, OsSMAX1 levels remained unaffected following treatment with either 1 µM or 10 µM dMGer treatment, while OsSMAX1 levels were reduced by 10 µM (−)-GR24 treatment (Figure 1G). These results suggest that dMGer promotes the degradation of OsSMAX1 in a *D14L*-dependent manner. Together, these findings demonstrate that dMGer functions as an effective and selective KL analog in rice *in vivo*.

### dMGer binds to D14L and induces its structural changes *in vitro*

We then examined whether dMGer directly interacts with D14L. We first conducted dYLG (Supplementary Figure S1) assays. dYLG is hydrolyzed in the ligand-binding pocket of D14L/KAI2 and releases a fluorophore. In this assay, the test compounds and dYLG are co-incubated with recombinant D14L/KAI2. Test compounds that bind to the ligand-binding pocket of D14L/KAI2 reduce the fluorescence of the dYLG-derived fluorophore by competitively inhibiting the hydrolysis of dYLG. D14L was incubated with 1 µM dYLG and different concentrations of dMGer or (−)-GR24. dMGer significantly reduced dYLG hydrolysis at 1 µM and almost completely abolished it at 5–10 µM (Figure 2A). By contrast, (−)-GR24 did not affect dYLG hydrolysis even at 10 µM. These results suggest that dMGer, but not (−)-GR24, binds to the ligand binding pocket of D14L.

**Figure 2.**
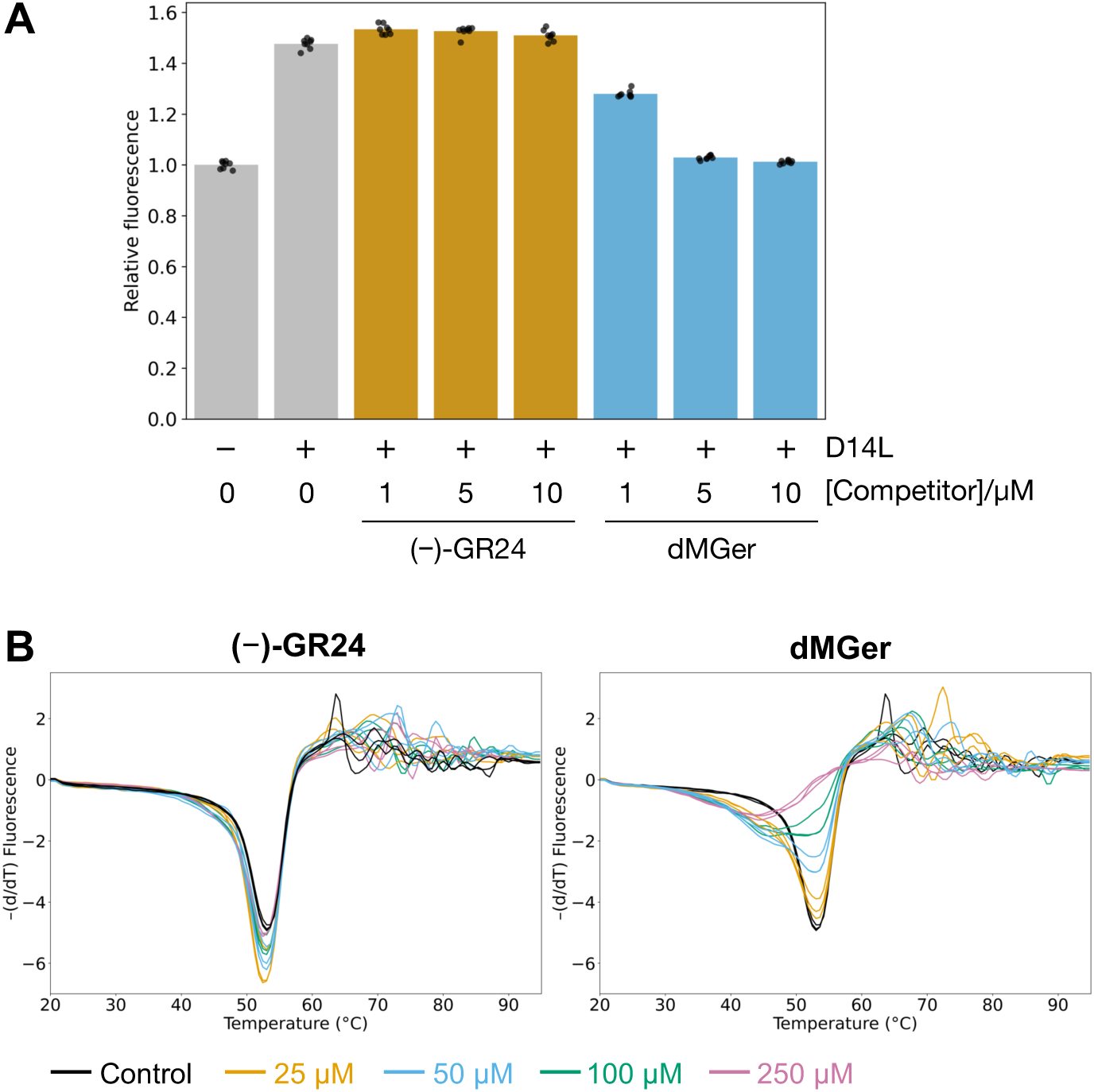
Binding of (−)-GR24 and desmethyl germinone to D14L *in vitro*. **(A)** Inhibitory effects of dMGer and (−)-GR24 on dYLG hydrolysis by MBP-6×His-D14L. Reactions contained 1 µM dYLG, 10 ng/µL protein, and the indicated concentrations of (−)-GR24 or dMGer. Dimethyl sulfoxide without either compound served as the mock control. Circles represent individual data points, expressed relative to the mean fluorescence intensity of mock reactions; bars indicate means (*n* = 8). **(B)** Differential scanning fluorimetry (DSF) of MBP-His-D14L in the presence of 0–100 µM (−)-GR24 (left) or desmethyl germinone (dMGer) (right). Melting curves of the protein incubated with the indicated concentrations of each compound are shown in different colors. Each line represents a single replicate (triplicates for each concentration). Control experiments, in which the protein was incubated without KL analogs, are shown in both plots. Alt text: Panels labeled A and B. A is a graph showing the competitive inhibition of D14L hydrolase activity by desmethyl germinone or (−)-GR24. B presents graphs showing the results of differential scanning fluorimetry in the presence of the KAI2 ligand analogs.

We also examined the binding of these compounds to D14L using DSF. D14L showed no Tm shift upon addition of (−)-GR24 (Figure 2B). In contrast, D14L showed a clear, dose-dependent decrease in Tm upon addition of dMGer (Figure 2B), indicating the ability of dMGer to directly interact with and change the structure of D14L.

### dMGer enhances the direct interactions of D14L with D3 and OsSMAX1

We next examined the interactions between the KL signaling components *in vitro*. We first performed pull-down assays using maltose-binding protein (MBP)-tagged D3 (MBP-D3) and Glutathione-S-transferase (GST)-tagged D14L. We found that MBP-D3, but not the control protein MBP-hexahistidine (His), interacted weakly with GST-D14L in the absence of KL analogs. This interaction was enhanced by dMGer, but not by (−)-GR24 (Figure 3A). We then performed pull-down assays using MBP-His-OsSMAX1 and GST-D14L. These proteins interacted even in the absence of KL analogs, and their interaction was also strengthened by the addition of dMGer (Figure 3B). In addition, the pull-down assay using MBP-D3 and GST-OsSMAX1 showed an interaction between these proteins in the absence of KL analogs (Figure 3C). These results demonstrate that the components of the KL signaling pathway are capable of directly interacting with one another *in vitro* and that dMGer enhances the interactions of D14L with D3 and OsSMAX1, suggesting its critical role in promoting the assembly of the KL signaling complex.

**Figure 3.**
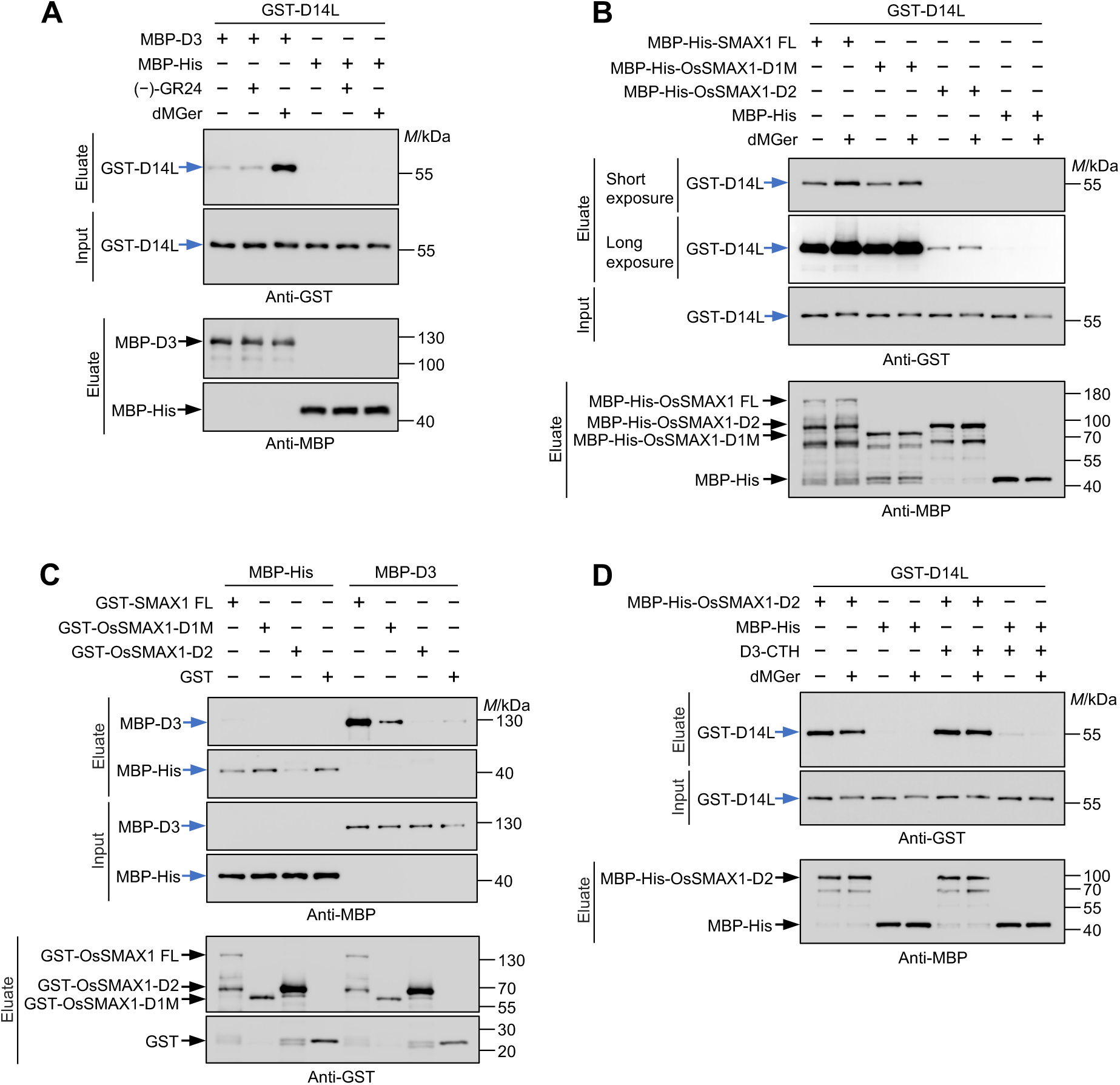
Interactions between KL signaling components. **(A)** *In vitro* MBP pull-down assay using dextrin agarose beads, with MBP-D3 co-expressed with ASK1 or MBP-His tag alone as bait, and GST-D14L as prey. All reactions contained 0.2% (v/v) DMSO as solvent for chemicals. Assays were supplemented with either 10 µM dMGer, 10 µM (−)-GR24, or DMSO only. Calculated molecular masses: GST-D14L, ∼56 kDa; MBP-D3, ∼121 kDa; MBP-His tag, ∼45 kDa. **(B)** *In vitro* MBP pull-down assay using MBP-His-tagged full-length (FL) or domain fragments of OsSMAX1 as bait, and GST-D14L as prey. Assays were performed with or without 10 µM dMGer. For better visualization of GST-D14L pulled-down by each bait, images of the same blot taken with different exposure times (short and long) are shown. Calculated molecular masses: MBP-His-OsSMAX1 FL, ∼154 kDa; MBP-His-OsSMAX1-D1M, ∼79 kDa; MBP-His-OsSMAX1-D2, ∼90 kDa. **(C)** *In vitro* GST pull-down assay using glutathione agarose beads, with GST-tagged FL or domains of OsSMAX1 as bait, and MBP-D3 or MBP-His as prey. Calculated molecular masses: GST-OsSMAX1 FL, ∼135 kDa; GST-OsSMAX1-D1M, ∼60 kDa; GST-OsSMAX1-D2, ∼71 kDa; GST tag, ∼28 kDa. **(D)** *In vitro* MBP pull-down assay using MBP-His-OsSMAX1-D2 as bait and GST-D14L as prey, performed in the presence or absence of D3-CTH peptide, with or without 10 µM dMGer. **(E)** GST- and MBP(-His)-tagged proteins were detected by immunoblotting with anti-GST and anti-MBP antibodies, respectively. The positions of size markers electrophoresed with the samples are shown alongside each blot. Alt text: Immunoblots labeled A to D showing physical interactions among D14L, D3, and OsSMAX1 proteins. A shows interaction between D14L and D3 promoted by desmethyl germinone. B shows interaction of D14L with full-length or D1M region of OsSMAX1 and effects of desmethyl germinone on the interactions. C shows interaction between D3 and OsSMAX1. D shows effect of D3-C-terminal helix on interaction between D14L and D2 domain of OsSMAX1.

### Domains of OsSMAX1 and D53 differentially interact with D14L or D14

As we demonstrated the interaction of D14L and OsSMAX1 and the enhancement of the interaction by dMGer *in vitro*, we next examined the functions of the domains of OsSMAX1. To compare the affinity of the OsSMAX1 D1M and D2 domains to D14L, we pulled down GST-D14 using MBP-His-OsSMAX1-D1M or MBP-His-OsSMAX1-D2. The D1M domain showed a stronger interaction with D14L than the D2 domain. While dMGer enhanced the interaction of the D1M domain with D14L, it did not enhance the interaction of the D2 domain with D14L (Figure 3B). As it has been shown that D3-CTH enhances the interaction between the D53-D2 domain and D14 (Shabek et al., 2018), we examined whether D3-CTH enhances the interaction between the OsSMAX1-D2 domain and D14L. However, D3-CTH did not enhance the interaction between OsSMAX1-D2 domain and D14L either in the absence or presence of dMGer (Figure 3D). We also examined the interaction of the OsSMAX1 domains with D3. The pull-down assay showed an interaction between OsSMAX1-D1M and D3 (Figure 3C). These results demonstrate that the OsSMAX1-D1M domain plays a central role in the interaction of OsSMAX1 with D14L and D3.

As our results suggested that the functions of OsSMAX1 domains for the interaction with D14L are different from those of D53 domains for the interaction with D14, we investigated the interactions of OsSMAX1 and D53 domains with D14L and D14. In our *in vitro* pull-down assays, D14L strongly interacted with the OsSMAX1-D1M domain, while it hardly interacted with the D53-D1M domain in the presence of dMGer. Although OsSMAX1-D2 and D53-D2 domains seem to have similar levels of affinity for D14L, these interactions were much weaker than the interaction between D14L and the OsSMAX1-D1M domain (Figure 4A). In contrast, D14 showed a stronger interaction with the D53-D2 domain in the presence of the D14 ligand (+)-GR24 than with the interactions with D53-D1M, OsSMAX1-D1M, and OsSMAX1-D2 domains (Figure 4B). This D14–D53-D2 interaction was enhanced by D3-CTH in a manner consistent with the previous report (Shabek et al., 2018). D3-CTH also increased the interaction efficiency between D14 and the OsSMAX1-D2 domain. The affinities of D14 with the two D2 domains in the presence of D3-CTH did not show any apparent difference (Figure 4B). These results suggest that D14–D53 and D14L–OsSMAX1 complexes exhibit distinct binding modes.

**Figure 4.**
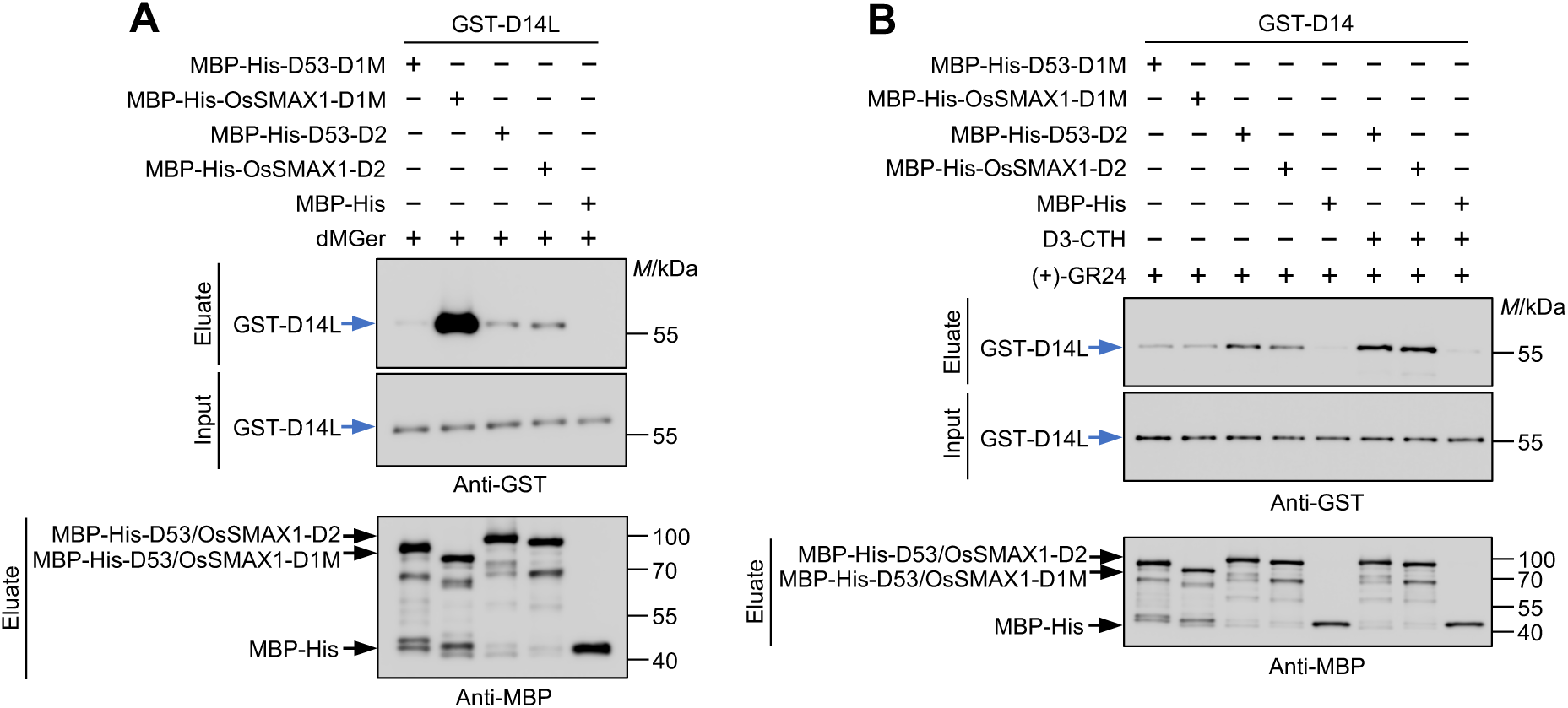
Differential interaction of D14L and D14 with domains of OsSMAX1 and D53. **(A)** *In vitro* MBP pull-down assay using MBP-His-tagged domains of D53 or OsSMAX1 as bait, and GST-D14L as prey. All reactions contained 10 µM dMGer. Calculated molecular masses: MBP-His-D53-D1M, ∼80 kDa; MBP-His-D53-D2, ∼91 kDa. **(B)** *In vitro* MBP pull-down assay using MBP-His-tagged domains of D53 or OsSMAX1 as bait, and GST-D14 as prey, performed in the presence or absence of D3-CTH. All reactions contained 10 µM (+)-GR24. Calculated molecular masses: GST-D14 ∼56 kDa. GST- and MBP-His-tagged proteins were detected by immunoblotting with anti-GST and anti-MBP antibodies, respectively. The positions of size markers electrophoresed with the samples are shown alongside each blot. Alt text: Immunoblots labeled A and B showing selective interactions between components of the KAI2 ligand and strigolactone signaling pathway. A presents interactions of D14L with domains of OsSMAX1 and D53, showing a stronger binding to D1M of OsSMAX1. B presents interactions of D14 with the same protein domains, showing a higher interaction with D2 of D53 enhanced by D3-C-terminal helix peptide.

## Discussion

Although the model of the KL signaling pathway, in which its components interact in a ligand-dependent manner, is widely accepted, the ligand-dependent interactions among these components have not been fully demonstrated. Using the recently developed artificial KL analog, dMGer, we addressed this issue. We show that dMGer effectively and selectively induces KL responses in rice and directly interacts with D14L. Furthermore, we demonstrate that the direct interaction between D3 and D14L is induced by dMGer, but not by (−)-GR24. We also report, for the first time, the enhancement of the interaction between D14L/KAI2 and SMAX1 by KL analogs *in vitro*. These findings demonstrate that these three proteins directly interact in a KL-responsive manner. Furthermore, parallel comparisons of interactions of D14L and D14 with domains of SMXL and D53 revealed regions that contribute to the specificity of KL and SL signaling. Collectively, our findings advance the understanding of the KL signaling by addressing knowledge gaps regarding the ligand perception and the interactions among the signaling components.

This work is the first report describing the physiological responses of rice to desmethyl-type KL analogs. dMGer has been shown to suppress hypocotyl elongation in a KAI2-dependent manner and to induce seed germination; however, evaluation of its physiological activities has been largely limited to Arabidopsis, except for germination-induction assays with the root parasitic plants *Orobanche minor* and *Striga hermonthica*, in which KAI2-dependency was not examined (Okabe et al., 2023). In the case of other desmethyl-type analogs, their physiological activity has been investigated only in Arabidopsis and *Brachypodium distachyon* (Yao et al., 2021; Meng et al., 2022). The observed D14L-dependent effects on plant growth, the regulation of KL marker genes, and the protein levels of OsSMAX1 demonstrate the function of dMGer as a KL analog in rice, suggesting the potential of desmethyl-type analogs in a wide range of species.

In our *in vitro* assays, dMGer and (−)-GR24 exhibited distinct activities. dMGer inhibited dYLG hydrolysis by D14L and induced a Tm shift of D14L in DSF assays, indicating direct binding to the D14L ligand-binding pocket and induction of structural changes in D14L, whereas (−)-GR24 did not induce these responses. Consistently, dMGer, but not (−)-GR24, enhanced the interaction between D14L and D3. Similarly, in Arabidopsis, although (−)-GR24 induces a slight Tm shift of KAI2 in DSF assay, the shift is much weaker than that induced by a desmethyl-type KL analog dGR24 (Yao et al., 2021). (−)-GR24 only marginally enhanced the interaction between KAI2 and MAX2 *in vitro* (Xu et al., 2018). These results imply that the *in vitro* activity of (−)-GR24 toward Arabidopsis KAI2 is also limited. The use of desmethyl-type KL analogs may also reveal the interaction of the KL signaling components in Arabidopsis.

Despite the lack of *in vitro* activity of (−)-GR24 toward D14L, it induces KL responses in rice. Several possible explanations could account for the discrepancy between the *in vitro* and *in vivo* activities of (−)-GR24. One explanation is the requirement for *in vivo* conversion of (−)-GR24 into an active molecule, as suggested in the case of KARs. Previous studies demonstrated that neither KAR_1_ nor KAR_2_ shifts the Tm of KAI2 in DSF assays, and that KAR_1_ is slow to induce ubiquitination of SMXL2 (Waters et al., 2015; Wang et al., 2020b). These findings suggest that metabolic processing renders KARs active. (−)-GR24 may likewise undergo metabolic conversion into an unidentified bioactive form *in vivo*. An alternative hypothesis is that KAI2/D14L is subject to post-translational modifications that alter its mode of ligand recognition. Various enzymatic and non-enzymatic post-translational modifications that influence catalytic activity or protein folding have been documented (Kokkinidis et al., 2020). Another possibility is that D14L recognizes (−)-GR24 only in the presence of plant-derived cofactors. As an example of cofactor requirements in plant hormone signaling, binding of inositol pentakisphosphate is essential for CORONATINE INSENSITIVE 1 and JAZ proteins to function as the co-receptor of jasmonate (Sheard et al., 2010).

Taking advantage of the *in vitro* activity of dMGer, we characterized the interaction mode of the KL signaling components. In the absence of dMGer, although the interaction between D3 and D14L was hardly detectable, both D3 and D14L interacted with OsSMAX1, indicating that D14L, D3, and OsSMAX1 can form a complex even in the absence of the ligand. In the presence of dMGer, the interactions of D14L with D3 and OsSMAX1 were enhanced. These results suggest that the function of KL is not to assemble three spatially separated components into a single complex, but rather to modulate interactions within the complex, presumably inducing conformational changes required for the ubiquitination of OsSMAX1, although further *in vivo* evaluation is required to confirm this model (Figure 5A).

**Figure 5.**
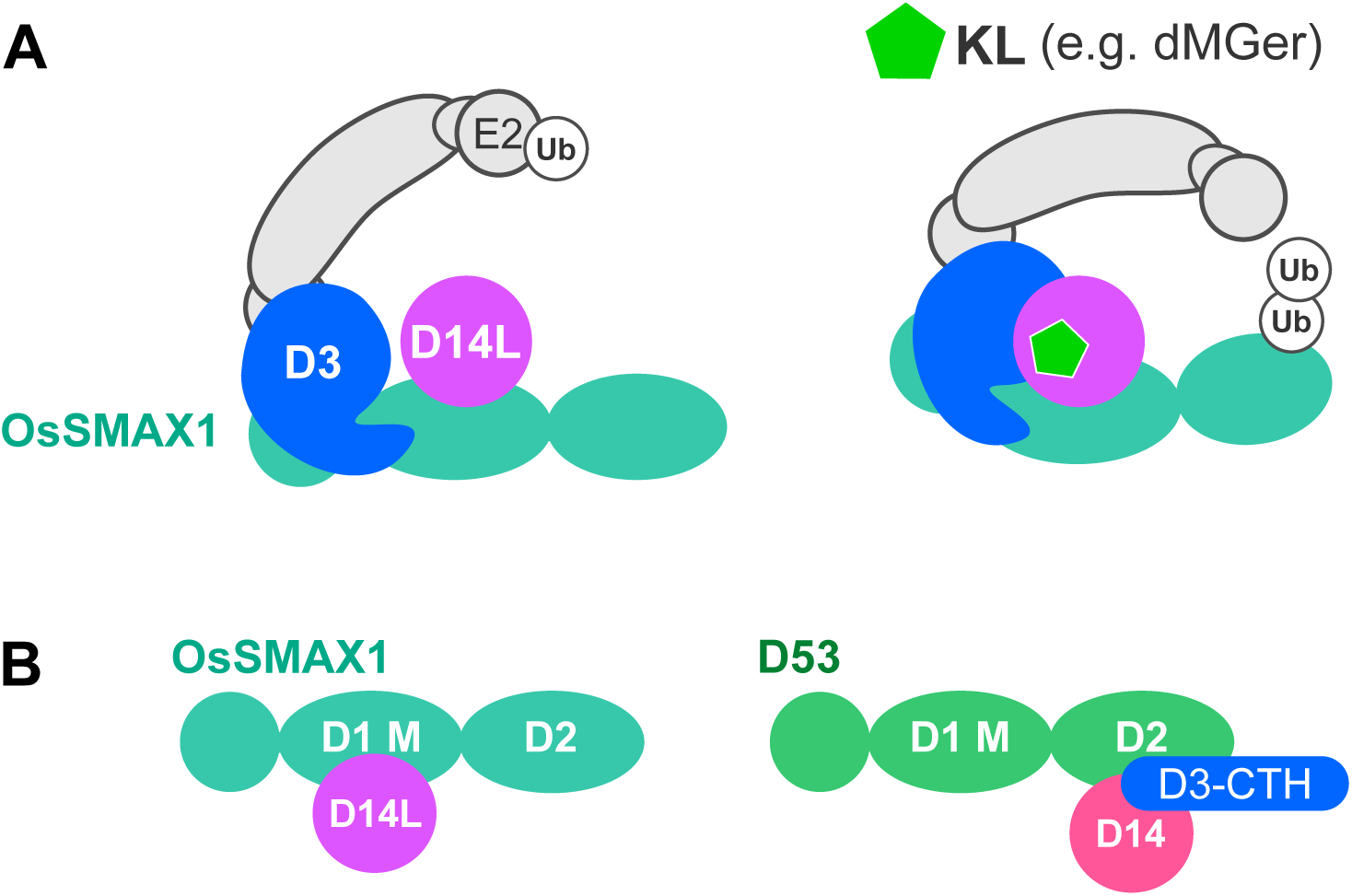
Interactions between rice KL signaling components and effects of KL. **(A)** Model of KL-triggered changes in protein–protein interactions. Even in the absence of KL, OsSMAX1 interacts with both D14L and D3, potentially forming a single complex. KL promotes direct binding between D14L and D3 and enhances the interaction between D14L and OsSMAX1, likely stabilizing and altering the conformation of the complex, thereby leading to the ubiquitination of OsSMAX1. Differences in interaction strength are indicated by the thickness of grey lines. **(B)** Schematic of physical interactions of rice D14L and D14 with domains of OsSMAX1 and D53. D14L mainly interacts with OsSMAX1-D1M, whereas D14 mainly interacts with D53-D2. D3-CTH enhances the interaction between D14 and the D53-D2 domain. Differences in interaction strength are indicated by the thickness of grey lines. Alt text: Graphical abstracts of models derived from study results, labeled A and B. A illustrates how ligand perception by D14L leads to protein–protein interactions and signal transduction. B illustrates differences in the interactions between receptors and suppressor proteins in the KAI2 ligand and strigolactone signaling pathways.

Furthermore, we clarified differences in the mode of interaction between the KL and SL signaling complexes. The selective interaction of D14L with OsSMAX1 relies on its strong affinity to the OsSMAX1-D1M domain rather than the D2 domain. On the other hand, D14 prefers the D53-D2 domain rather than the D1M domain. D3-CTH enhances the interaction between D14 and D53-D2, but not the interaction between D14L and OsSMAX1-D2 (Figure 5B). Previous studies have shown that D3/MAX2-CTH is involved in the KL signaling (Tal et al., 2022; Tal et al., 2023). Our results suggest that D3/MAX2-CTH functions in the KL signaling pathway are different from those in the SL signaling. These findings demonstrate that although D14 and D14L, as well as D53 and OsSMAX1, are paralogs, they distinctively interact in the KL and SL signaling complexes. Intriguingly, D3-CTH also enhanced the interaction of D14 with OsSMAX1-D2 under our assay conditions. Previously, the *in vivo* interaction between AtD14 and SMAX1/SMXL2 has been reported (Wang et al., 2020b; Li et al., 2022). This interaction may be mediated by the SMAX1-D2 domain and D3/MAX2-CTH. Further biochemical and structural dissection of SMAX1 and D53 domains would elucidate the molecular basis underlying the divergence of the KL and SL signaling pathways.

## Materials and Methods

### Plant materials and growth conditions

The rice cultivar Nipponbare was used as the wild type. The *d14l-c1* line was previously described (Mashiguchi et al., 2023). The *d14l-c2* line was generated using the same gRNA as that for *d14l-c1*, as described previously (Mashiguchi et al., 2023). Due to the duplication of the 174th cytosine in the D14L ORF, a frame shift occurs after the 59th amino acid residue, generating a premature stop codon. The previously described *d14l-1* line in the Nipponbare genetic background (Zheng et al., 2020) was used for the protein degradation assay.

To test mesocotyl responses to dMGer, rice seeds were surface-sterilized with sodium hypochlorite for 15 minutes and placed on 0.6% (w/v) agarose plates supplemented with the indicated concentrations of the compounds. A 1000× stock solution of dMGer, dissolved in acetone, was added to the medium to achieve the final concentrations. Plates containing 0.1% acetone served as mock controls. Seeds were germinated and grown in the dark at 28 °C for 8 days in an incubator. To remove seedlings that did not germinate properly from the analysis, only seedlings that properly developed embryonic roots and had a ≥2-cm of aerial part were measured. Mesocotyl length was measured using Fiji.

To test the effects of chemicals on *DLK2b* and *KUF1* transcription, rice seeds were surface-sterilized with sodium hypochlorite for 15 minutes and then grown on solid Hoagland medium containing 0.8% Phytagel for 4 days at 28 °C under a 12-h light/12-h dark cycle. Uniform seedlings were then selected and transferred to 1 mL of Hoagland solution, cultured overnight, and then incubated for 6 hours with 0.1, 1, or 10 µM of (−)-GR24 or dMGer at 28°C. A 1000× stock solution of each compound, dissolved in acetone, was used to achieve the final concentrations. Hoagland solution containing 0.1% (v/v) acetone without any added compound was used as the mock control.

Primary calli of WT and the *d14l-1* mutants (Zheng et al., 2020) for protein degradation assay were induced on solid induction medium at 28°C under a photoperiod of 22 hours light followed by 2 hours dark for a duration of 10 days (Jiang et al., 2013). Subsequently, the primary calli were transferred onto plates containing solid induction medium for subculturing. The calli were maintained on these plates for a maximum of 14 days, after which they required transfer to fresh plates to ensure continued growth.

### qRT-PCR analysis

Total RNA was extracted from rice roots using the EasyPure Plant RNA Kit (TransGen). An aliquot (0.05–0.2 µg) was used for first-strand cDNA synthesis with the TransScript Uni All-in-One First-Strand cDNA Synthesis SuperMix for qPCR (TransGen). RT-qPCR was performed using gene-specific primers (Supplementary Table S1) on an Agilent Mx3000P system, following the manufacturer’s instructions. Each 10 µL reaction contained 1 µL of diluted cDNA, 0.2 µM of each primer, and 5 µL of KOD SYBR qPCR Mix (Toyobo). The rice *UBQ* gene was used as an internal reference.

### Protein degradation assay

Twice sub-cultured calli were collected and divided into separate 10 ml tubes containing 3 ml fresh liquid induction medium, then incubated at 28°C on orbital shaker (80 rpm) for 2 hours. After this pre-incubation, dMGer, or (−)-GR24 at specified concentrations were added to the samples for treatment. Following the treatment for two hours, callus samples were quickly collected and immediately frozen in liquid nitrogen. The stock solution of dMGer and (−)-GR24 were prepared by dissolving them in acetone. Acetone without any added compound was used as the mock control.

For protein extraction, the frozen samples were ground into powder. Protein extraction buffer (50 mM Tris-HCl, pH 7.5, 150 mM NaCl, 5 mM EDTA, 1% Triton X-100, 0.5% (w/v) SDS, 0.5% β-mercaptoethanol, 50 μM MG132, 5 mM DTT, 1× protease inhibitor cocktail, and 1 mM PMSF) was added to the powdered callus tissue. The mixture was vortexed for 30 seconds and then incubated on ice for 2 minutes; this process was repeated five times to ensure complete cell lysis. The lysate was centrifuged at 12,000 rpm for 10 minutes at 4°C, and the supernatant was collected. Subsequently, 5×SDS loading buffer was added to the supernatant, which was then heated at 95°C for 10 minutes to denature the proteins.

Proteins were separated by SDS-PAGE and transferred onto PVDF membranes. Detection of OsSMAX1 was performed as described by Zheng et al. (2020), using polyclonal antibodies against OsSMAX1 at a 1:5000 dilution. ACTIN was used as internal loading controls and detected with anti-ACTIN (1:5000) antibody (Abclonal).

### Plasmid construction

To construct the plasmid for expressing His-D14L for DSF assays, the ORF fragment of *D14L* was amplified by PCR from cDNA synthesized from rice total RNA. The PCR product was cloned by In-Fusion (Takarabio) into the pET47b vector (Novagen) using SmaⅠ and SacⅠ sites. To construct the plasmid for expressing MBP-His-D14L, the CDS of *D14L* was amplified from cDNA synthesized from rice total RNA, and the PCR product was inserted into the Sma1 and EcoR1 sites of the modified pMAL-c5x vector (Seto et al., 2019) (pMALHis). For expression of GST-D14L, the *D14L* CDS was subcloned from pMALHis-D14L into pGEX-4T-3 (Cytiva) using EcoRI and NotI sites. For expression of GST-D14, the *D14* (50–318) CDS in cDNA synthesized from rice total RNA was amplified and inserted into pGEX-4T-3 using EcoRI and NotI sites.

For the bacterial expression of MBP-His-tagged full-length OsSMAX1 (residues 2–1041) (OsSMAX1-FL) and domains of OsSMAX1 (D1M: 218–541; D2: 638–1041), the corresponding CDS regions were amplified from cDNA synthesized from rice total RNA. PCR products were first inserted into pGEX-4T-3, followed by subcloning into pMALHis, replacing D14L in pMALHis-D14L.

For the bacterial expression of MBP-His-tagged domains of D53, the fragments encoding the D1M (182–510) and D2 (718–1131) domains of D53 were amplified from rice genomic DNA. The D53-D1M fragment was produced by overlap extension PCR using two fragments amplified from exon 1 and exon 2 of genomic *D53*. A nonsynonymous substitution found in the amplified D53-D1M fragment was corrected by overlap extension PCR. These fragments were cloned into pMALHis.

To construct a plasmid for co-expression of MBP-tagged D3 (2–720) and untagged ASK1 in insect cells, the CDSs of *D3* and *ASK1* were amplified from cDNA synthesized from total RNA extracted from rice and Arabidopsis seedlings, respectively. ASK1 was inserted into pFastBac Dual (Thermo Fisher Scientific) at the *Nco*I site downstream of the p10 promoter, yielding pFastBacDual-ASK1. *D3* was first cloned into pGEX-4T-3 and then subcloned into pFastBacDual-ASK1 downstream of the polyhedrin promoter, fused with an MBP tag amplified from pMALHis. For multi-fragment assembly, we used the ClonExpress II One Step Cloning Kit (Vazyme).

Primers used for the plasmid construction are listed in Supplementary Table S1.

### Preparation of recombinant proteins

*E. coli* BL21(DE3) was used for the expression of D14L for DSF. The overnight culture (20 mL) was added to a fresh LB medium (2.0 L) containing kanamycin (50 mg/L) at 37°C. After OD600 reached 0.8, 0.25 mM isopropyl β-D-1-thiogalactopyranoside (IPTG) was added, and the culture was further incubated at 18 °C for 21 h. The culture medium was centrifuged at 3500 *g* and the pellet was stored at −30°C until use. The pellet was resuspended and sonicated in lysis buffer (50 mM Tris buffer (pH 8.0) containing 500 mM NaCl, 5 mM 2-mercaptoethanol, and 10 % glycerol). The supernatant was purified using His60 Ni Superflow Resin (1 mL, Takara). After washing with the washing buffer (50 mM Tris buffer (pH 8.0) containing 500 mM NaCl and 20 mM imidazole), the bound protein was eluted with elution buffer (50 mM Tris buffer (pH 8.0) containing 500 mM NaCl and 200 mM imidazole). The eluate was concentrated using VIVASPIN Turbo 15 (Sartorius), adjusted to 12 mg/mL, aliquoted, immediately frozen in liquid nitrogen, and stored at −80°C until use.

*E.coli* strain Rosetta (DE3) (Novagen) was used for the production of recombinant proteins, except for D3 and ASK1, for *in vitro* pull-down assays and dYLG assays. Transformed cells were cultured in LB media containing 50 µg/mL carbenicillin and 50 µg/mL chloramphenicol at 16 °C until the OD₆₀₀ reached 0.4–0.6. Protein expression was induced with 0.1 mM IPTG, and cells were incubated overnight at 16 °C.

For purification of MBP-6×His-fused D14L, OsSMAX1-D1M, OsSMAX1-D2, OsSMAX1-FL, D53-D1M, D53-D2, and the MBP-6×His control protein, cells were harvested by centrifugation and lysed by sonication in lysis buffer (25 mM Tris-HCl, pH 8.0, 500 mM NaCl, 1 mM TCEP, 20 mM imidazole, and 0.4 mM phenylmethylsulfonyl fluoride (PMSF)). The lysate was centrifuged at 18,000 rpm for 50 min at 4 °C. The supernatant was applied to Ni NTA Beads (Smart-Lifesciences) in a Poly-Prep Chromatography Column (Bio-Rad). After washing with lysis buffer, proteins were eluted with MBP elution buffer (20 mM Tris-HCl, pH 8.0, 500 mM NaCl, 1 mM TCEP, and 250 mM imidazole). The eluate was subsequently applied to Dextrin Beads (Smart-Lifesciences) in a Poly-Prep Chromatography Column. After washing with MBP wash buffer (20 mM Tris-HCl, pH 8.0, 500 mM NaCl, 1 mM TCEP), proteins were eluted with MBP elution buffer (MBP wash buffer containing 10 mM maltose). Eluted proteins were concentrated and subjected to buffer exchange using Amicon Ultra-15 (10 kDa MWCO for MBP-6×His, 30 kDa for the others). The buffer was exchanged first to MBP wash buffer, and then to either 50 mM Tris-HCl, pH 7.6, 150 mM NaCl, 0.5 mM TCEP (for D14L), 20 mM Tris-HCl, pH 7.5, 500 mM NaCl, 0.5 mM TCEP (for OsSMAX1-D1M, and D53-D1M), 50 mM Tris-HCl, pH 7.5, 500 mM NaCl, 0.5 mM TCEP (for OsSMAX1-D2 and D53-D2), or 20 mM Tris, pH 8.0, 500 mM NaCl, 0.5 mM TCEP (for OsSMAX1-FL, MBP-6xHis). In the final dialyzed samples, the residual concentrations of MBP elution and MBP wash buffers were less than 0.2% and 1% (v/v), respectively.

For purification of GST-D14L and GST-D14, cells were harvested by centrifugation and lysed by sonication in lysis buffer (25 mM Tris-HCl, pH 7.6, 500 mM NaCl, 1 mM TCEP, and 0.4 mM PMSF). The lysate was centrifuged at 18,000 rpm for 50 min at 4 °C. The supernatant was applied to Glutathione Beads (Smart-Lifesciences) packed in a Poly-Prep Chromatography Column. After sequential washing with lysis buffer and GST wash buffer (100 mM Tris-HCl, pH 8.0, 500 mM NaCl, 1 mM TCEP), proteins were eluted with GST elution buffer (100 mM Tris-HCl, pH 8.0, 500 mM NaCl, and 20 mM reduced glutathione). Eluted proteins were concentrated and subjected to buffer exchange using Amicon Ultra-15 centrifugal filters (30 kDa MWCO). The buffer was first exchanged to dialysis buffer 1 (100 mM Tris-HCl, pH 8.0, 500 mM NaCl, 0.5 mM TCEP), and then to 50 mM Tris-HCl, pH 7.5, 150 mM NaCl, 0.5 mM TCEP. In the final dialyzed samples, the residual concentrations of GST elution and dialysis buffer 1 were less than 0.2% and 1% (v/v), respectively.

For purification of GST-fused OsSMAX1-D1M, OsSMAX1-D2, and OsSMAX1-FL, the same lysis, GST wash, and GST elution buffers as those for the purification of GST-D14L and GST-D14 were used. For OsSMAX1-D1M and OsSMAX1-FL, the sonicated cell lysate was mixed with polyethylenimine (Sinopharm Chemical Reagent Co., Ltd) to a final concentration of approximately 0.1%(w/v) before centrifugation at 18,000 rpm. Subsequent purification with Glutathione Beads was similar to that for GST-D14L/D14. Eluted proteins were concentrated and subjected to buffer exchange using Amicon Ultra-15 centrifugal filters (30 kDa MWCO). The buffer was first exchanged to dialysis buffer 1, and then to either 50 mM HEPES-NaOH, pH 7.0, 500 mM NaCl, 0.5 mM TCEP (for D1M and FL) or 50 mM HEPES-NaOH, pH 7.0, 500 mM NaCl, 1 mM TCEP (for D2).

MBP-tagged D3 was co-expressed with untagged ASK1 in *Spodoptera frugiperda* Sf9 cells. Suspension cultures were infected with recombinant baculovirus generated using the Bac-to-Bac Baculovirus Expression System (Thermo Fisher Scientific), following the manufacturer’s instructions. Cells were harvested by centrifugation and lysed by sonication in 50 mM Tris-HCl, pH 7.5, 200 mM NaCl, 0.5 mM TCEP, 1% Protease Inhibitor Cocktail (EDTA-Free) (APExBIO, K1007). The lysate was clarified by centrifugation at 18,000 rpm for 50 min at 4 °C, and the supernatant was applied to Dextrin Beads (Smart Lifesciences) in a Poly-Prep chromatography column (Bio-Rad). After washing with D3 wash buffer (50 mM Tris-HCl, pH 7.5, 200 mM NaCl, 0.5 mM TCEP), bound proteins were eluted with D3 elution buffer (D3 wash buffer supplemented with 10 mM maltose). Eluted proteins were concentrated and subjected to buffer exchange using Amicon Ultra-4 (10 kDa MWCO). The buffer was first exchanged to D3 wash buffer, followed by 50 mM Tris-HCl, pH 7.5, 150 mM NaCl, 0.5 mM TCEP. In the final dialyzed samples, the residual concentrations of D3 elution and D3 wash buffers were less than 0.1% and 1% (v/v), respectively.

### Differential scanning fluorometry experiments

DSF was conducted using 4.8 µg of the recombinant D14L protein in 20 µL of PBS buffer containing the 0.005 µL of Sypro Orange (Ex/Em: 490/610 nm; Invitrogen) and the tested compound with 5% (v/v) acetone on a 96-well plate. Mixtures were heated from 20°C to 95°C, and the fluorescence (Ex/Em; 490/610) was continuously scanned by using LightCycler480. The denaturation curve was calculated by using the LightCycler480 Software.

### dYLG hydrolysis assays

Reactions contained 1 µM dYLG, 0–5 µM competitor, 0.3% DMSO, and 10 µg/mL MBP-6×His-D14L. Recombinant protein (10 mg/mL in 50 mM Tris-HCl, pH 7.6, 150 mM NaCl, 0.5 mM TCEP) was diluted 500-fold with reaction buffer (100 mM HEPES, pH 7.0, 150 mM NaCl). dYLG (1 mM stock in DMSO) and the competitor (500× stock in DMSO) were diluted with reaction buffer to twice their final concentrations before addition to the reaction. Protein solution (25 µL/well) was dispensed into a 1/2 Area OptiPlate-96 (PerkinElmer), followed by the addition of an equal volume of the dYLG/competitor solution, on ice. Fluorescence intensity was measured at 30 °C in the dark (Ex/Et: 480/520 nm) using Varioskan Flash (Thermo Fisher Scientific).

### Pull-down assays

The of D14L with D3 or domains of OsSMAX1/D53 were assessed using 2.5 µg GST-D14L (prey protein) was incubated with an approximately equimolar MBP-His (as a control) or MBP-tagged bait proteins at 4 °C in 700 µL of incubation buffer (20 mM Bis-Tris-HCl, pH 7.0, 150 mM NaCl, 0.1% Triton X-100, 0.2% DMSO for D3; 20 mM Tris-HCl, pH 7.5, 150 mM NaCl, 0.1% Triton X-100, 0.2% DMSO for the others). After ∼1 h of incubation, 25 µL of 50% Dextrin Beads (Smart Lifesciences) slurry, equilibrated and suspended in incubation buffer without KL analog, were added and incubated at 4 °C for ∼1.5 h. Beads were washed three times with incubation buffer, and bound proteins were eluted with elution buffer (20 mM Bis-Tris-HCl, pH 7.0, 150 mM NaCl, 10 mM maltose for D3; 20 mM Tris-HCl, pH 7.5, 150 mM NaCl, 10 mM maltose for the others). Eluates were mixed with an equal volume of 2× SDS-PAGE sample buffer and boiled at 95 °C for 5 min. To test the effects of dMGer or (−)-GR24, 10 µM of each chemical was added to the incubation buffer. When testing the effects of D3-CTH, synthesized D3-CTH peptideswere added at an approximately equimolar to the prey protein, and the beads were washed with incubation buffer containing no peptide.

The interactions of D14 with domains of OsSMAX1/D53 were assessed using similar procedures to those for D14L. 10 µM (+)-GR24 was added to the incubation buffer (20 mM Tris-HCl, pH 7.5, 150 mM NaCl, 0.1% Triton X-100, 0.2% DMSO).

To assess the interactions between D3 and domains of OsSMAX1, 5 µg of MBP-D3-ASK1 and 2 µg of MBP-6×His (control) were incubated with an approximately equimolar amounts of GST-tagged bait proteins or GST (control) in 700 µL of incubation buffer (20 mM Tris-HCl, pH 7.5, 150 mM NaCl, 0.5 mM TCEP, 0.1% Triton X-100). After ∼1 h of incubation, 25 µL of 50% Glutathione Beads (Smart Lifesciences) slurry, equilibrated and suspended in incubation buffer, was added and incubated at 4 °C for ∼1.5 h. Beads were washed three times with incubation buffer, and bound proteins were eluted with elution buffer (100 mM Tris-HCl, pH8.0, 150 mM NaCl, 20 mM Reduced glutathione). Eluates were mixed with an equal volume of 2× SDS-PAGE sample buffer and boiled at 95 °C for 5 min.

Input and eluted proteins were resolved by SDS-PAGE and detected by western blotting. Western blots were performed using mouse anti-GST antibody (Beijing Protein Innovation, AbM59001-2H5-PU; 1:3000–1:1000 diluted), mouse anti-MBP antibody (Beijing Protein Innovation, AbM59007-3-PU; 1:6000–1:1000 diluted), and HRP-conjugated goat anti-mouse IgG (H+L) (ABclonal, AS003; 1:10000 diluted).

## Author contributions

H.K. and K.T. designed the research. K.T., J.W., Q.X., Y.H., T.S., Y.Y., Y.S., and G.X. performed research. K.T., J.W., Q.X, Y.H., T.S., Y.S., G.X., and H.K. analyzed data. K.T. and H.K. wrote the manuscript.

## Acknowledgements

We thank to Miaolian Ma (Eukaryotic Protein Expression and Purification Platform, CAS CEMPS) for technical support in protein expression using insect cell culture, Liu Li (Core Facility Center of CEMPS, CAS) for technical instruction on fluorescence measurements, Prof. Dr. Minrui Fan (CAS CEMPS) for access to the equipment for protein extraction, Prof. Dr. Ruiqiang Ye (CAS CEMPS) for access to the equipment for western blot, and Dr. Kosuke Fukui (Tokyo University of Science) for providing dMGer.

## Supplementary data

**Supplementary Figure S1. Structure of compounds used in this study.**

Chemical structures of desmethyl germinone (dMGer) (top left), desmethyl Yoshimulactone Green (dYLG) (top right), (−)-GR24 (bottom left), and (+)-GR24 (bottom right).

**Supplementary Figure S2. Uncropped blots.**

(**A**) Full western blot images of Figure 3A.

(**B**) Full western blot images of Figure 3B.

(**C**) Full western blot images of Figure 3C.

(**D**) Full western blot images of Figure 3D.

(**E**) Full western blot images of Figure 4A.

(**F**) Full western blot images of Figure 4B.

**Supplementary Table S1. List of primers used in this study.**

