## Supplementary Figure S1; Supplementary Figure S2; Supplementary Table S1 for "*In vitro* evaluation of protein–protein interactions in the rice KAI2 ligand signaling complex"

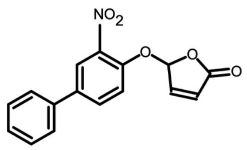

dMGer

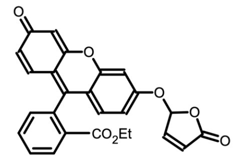

dYLG

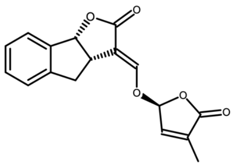

(−)-GR24

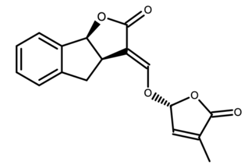

(+)-GR24

**Supplementary Figure S1. Structure of compounds used in this study.**

Chemical structures of desmethyl germinone (dMGer) (top left), desmethyl Yoshimulactone Green (dYLG) (top right), (−)-GR24 (bottom left), and (+)-GR24 (bottom right).

A

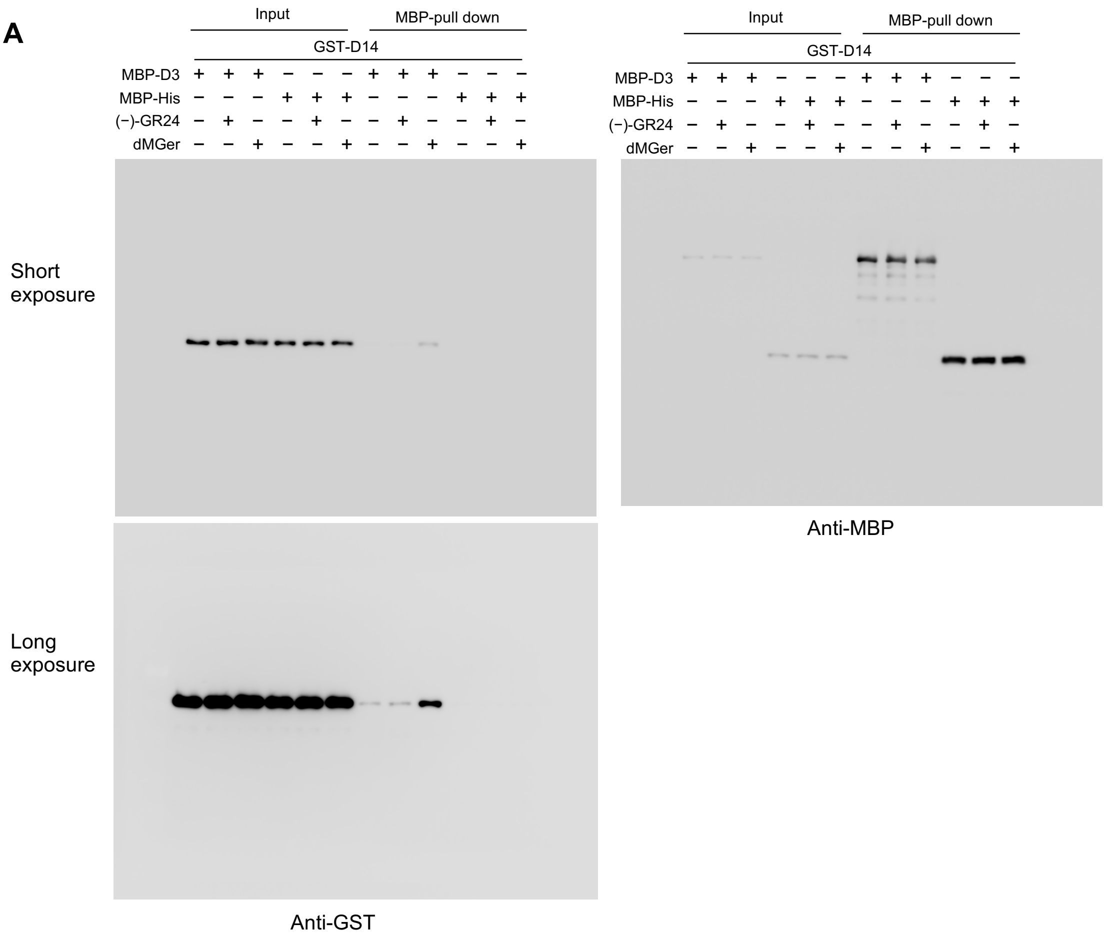
Input
MBP-pull down
GST-D14
MBP-D3
MBP-His
(−)-GR24
dMGer
Short exposure
Long exposure
Anti-GST
Anti-MBP

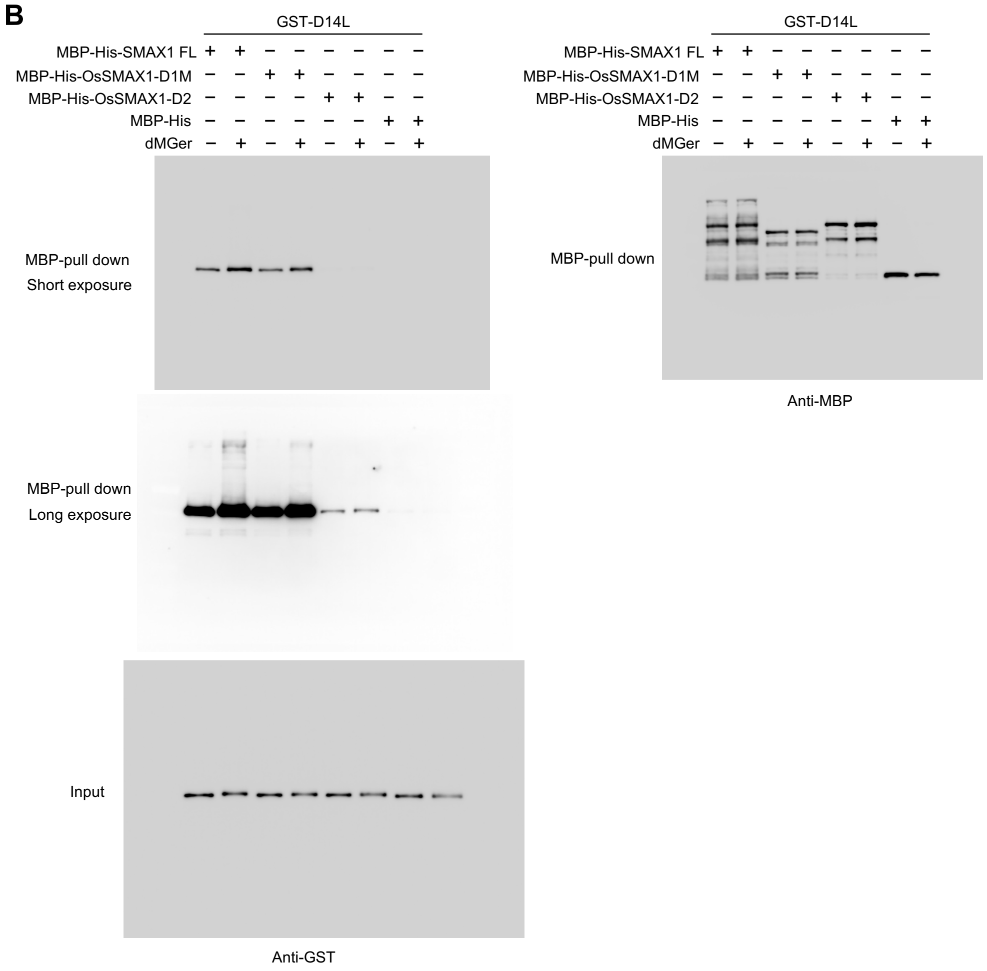
B
GST-D14L
MBP-His-SMAX1 FL + + − − − − − −
MBP-His-OsSMAX1-D1M − − + + − − − −
MBP-His-OsSMAX1-D2 − − − − + + − −
MBP-His − − − − − − + +
dMGer − + − + − + − +
MBP-pull down
Short exposure
MBP-pull down
Long exposure
Input
Anti-GST
GST-D14L
MBP-His-SMAX1 FL + + − − − − − −
MBP-His-OsSMAX1-D1M − − + + − − − −
MBP-His-OsSMAX1-D2 − − − − + + − −
MBP-His − − − − − − + +
dMGer − + − + − + − +
MBP-pull down
Anti-MBP

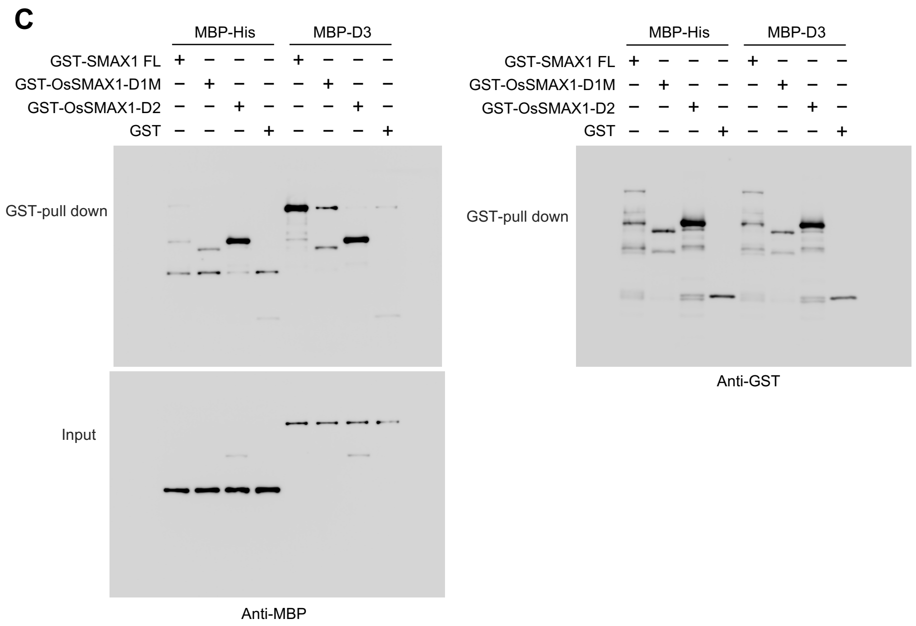

C
MBP-His
MBP-D3
GST-SMAX1 FL
GST-OsSMAX1-D1M
GST-OsSMAX1-D2
GST
GST-pull down
Input
Anti-MBP
Anti-GST


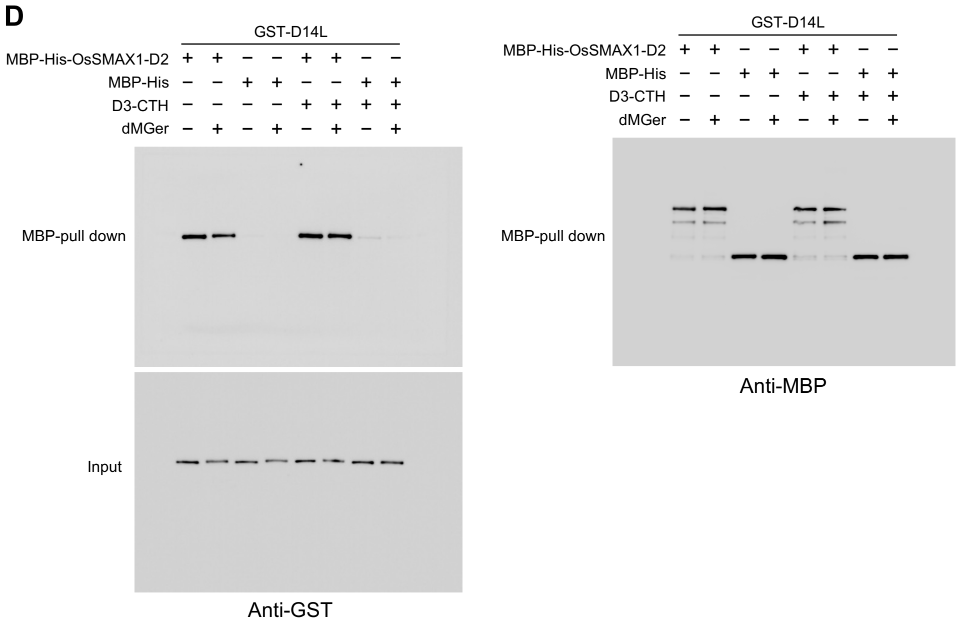

D
GST-D14L
MBP-His-OsSMAX1-D2
MBP-His
D3-CTH
dMGer
MBP-pull down
Input
Anti-GST
Anti-MBP

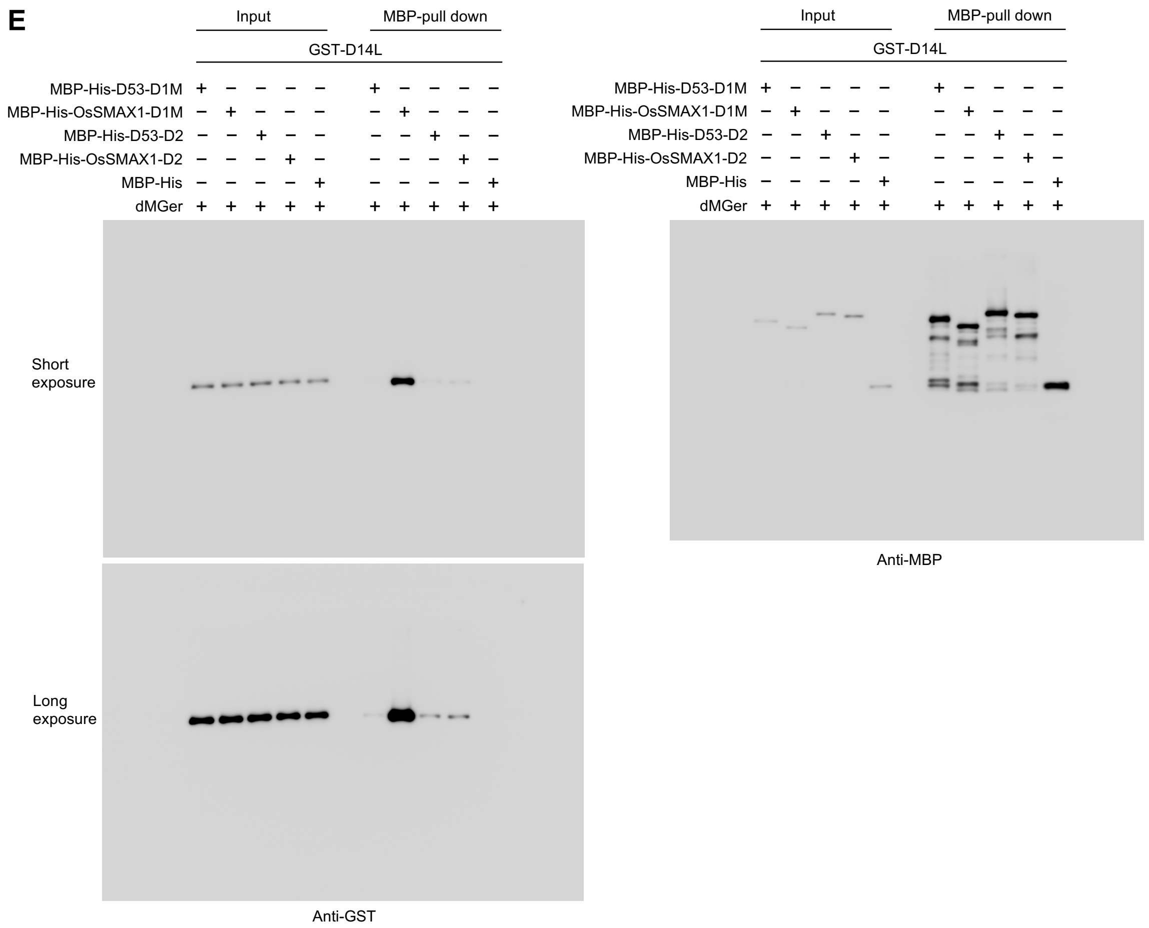
E
Input
MBP-pull down
GST-D14L
MBP-His-D53-D1M
MBP-His-OsSMAX1-D1M
MBP-His-D53-D2
MBP-His-OsSMAX1-D2
MBP-His
dMGer
Short exposure
Long exposure
Anti-GST
Input
MBP-pull down
GST-D14L
MBP-His-D53-D1M
MBP-His-OsSMAX1-D1M
MBP-His-D53-D2
MBP-His-OsSMAX1-D2
MBP-His
dMGer
Anti-MBP

F

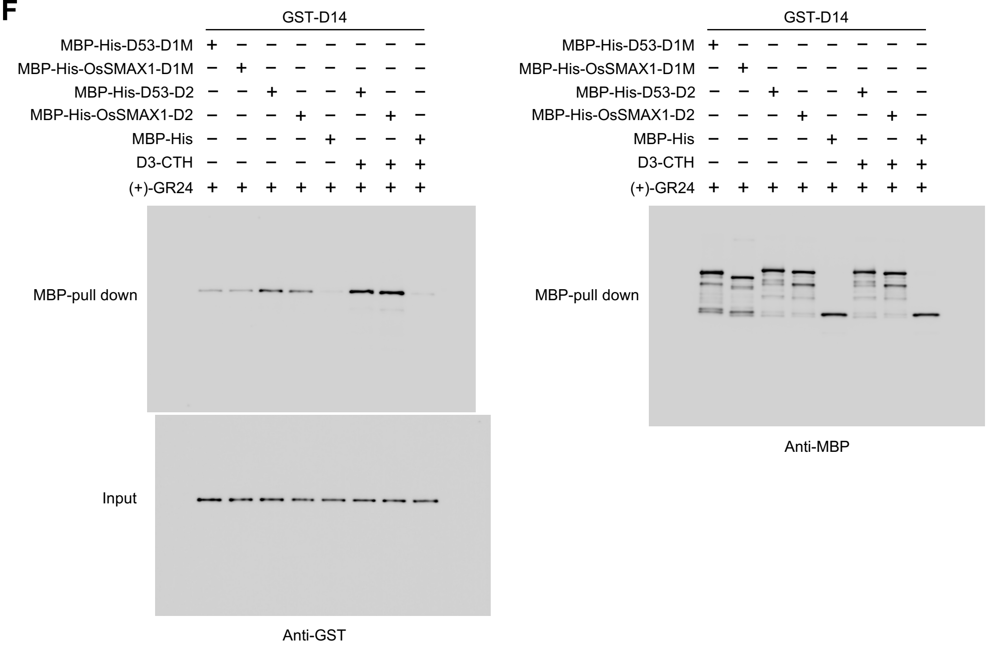


**Supplementary Figure S2. Uncropped blots.**

(**A**) Full western blot images of Figure 3A.

(**B**) Full western blot images of Figure 3B.

(**C**) Full western blot images of Figure 3C.

(**D**) Full western blot images of Figure 3D.

(**E**) Full western blot images of Figure 4A.

(**F**) Full western blot images of Figure 4B.

**Supplementary Table S1. List of primers used in this study.**

| Objective | Primer name | Sequence (5’ -> 3’) |
|---|---|---|
| Plasmid construction | pET47b-D14L_F | CTCTTTCAGGGACCCATGGGGATCGTGGAGGAGG |
| | pET47b-D14L_R | CACCAGAGCGAGCTCCTAGACCGCAATGTCATGCTGG |
| | pMALHis-D14L_F | ATGGGGATCGTGGAGGAGGC |
| | pMALHis-D14L_R | TTTTGAATTCCTAGACCGCAATGTC |
| | KT121_PGEX-D14L_F | CTGGTTCCGCGTGGATCCCCGAATTCCATGGGGATCGTGGAGGA |
| | KT122_pGEX-D14L_R | ATCGTCAGTCAGTCACGATGCGGCCGCCTAGACCGCAATGTCATGCTG |
| | KT104_pMAL-D14_F | TGAAGTCCTCTTTCAGGGACCCgcgccgagcggggcgaagct |
| | KT105_pMAL-D14_R | CTTATTTAATTACCTGCAGGGAATTCttagtaccgggcgagagcgcgg |
| | KT106_pGEX-D14_F | CTGGTTCCGCGTGGATCCCCGAATTCCgcgccgagcggggcgaagct |
| | KT107_pGEX-D14_R | ATCGTCAGTCAGTCACGATGCGGCCGCttagtaccgggcgagagcgcgg |
| | HR_OsSMAX1deltaATG_F | TCTGGTTCCGCGTGGATCCAGGGCGGATCTTAGCACC |
| | HR_OsSMAX1_R | TCAGTCACGATGCGGCCGCCTACATGCCATCAATGGCG |
| | BamH1_OsSMAX1D2_newF | aaGGATCCGCGGACTAACGGCTCTACAAAAGG |
| | NotI_OsSMAX1D2_R | aaGCGGCCGCCTACATGCCATCAATGGCG |
| | KT67_pGX-SMAX1D1_F | AATCGGATCTGGTTCCGCGTGGATCCGGCGCGGCCAACGCATACCT |
| | KT68_pGX-SMAX1M_R1 | GATCGTCAGTCAGTCACGATGCGGCCGCctaGGCGGAGTGGATTCGG GCACA |
| | KT119_pMAL-SMAX1_F | TTGAAGTCCTCTTTCAGGGACCCAGGGCGGATCTTAGCACCAT |
| | KT120_pMAL-SMAX1_R | CTTATTTAATTACCTGCAGGGAATTCCTACATGCCATCAATGGCGATCG |
| | KT112_pFBD-ASK1_F | GATCACCCGGGATCTCGAGCCATGATGTCTGCGAAGAAGATTGTGTTG |
| | KT113_pFBD-ASK1_R | TATGCATCAGCTGCTAGCACCATGGTCATTCAAAAGCCCATTGGTTCTC |
| | KT117_pFBD_GST_F2 | TACCGTCCCACCATCGGGCGCGACAACATGTCCCCTATACTAGGTTATT GGA |
| | KT118_pFBD_D3_R | GCTCGTCGACGTAGGCCTTTGAATTCCTAATCATCAATTTGCCGGCTGT |
| | KT135_pMAL-SMAX1D1_F | TTGAAGTCCTCTTTCAGGGACCCGGCGCGGCCAACGCATACCT |
| | KT136_pMAL-SMAX1M_R | CTTATTTAATTACCTGCAGGGAATTCctaGGCGGAGTGGATTCGGGCAC A |
| | KT133_pMAL-S1D2_F | TTGAAGTCCTCTTTCAGGGACCCGGACTAACGGCTCTACAAAAGG |
| | KT134_pMAL-S1D2_R | CTTATTTAATTACCTGCAGGGAATTCCTACATGCCATCAATGGCGAT |
| | KT163_pMAL_D53D1_F | TTGAAGTCCTCTTTCAGGGACCCatgccgctcctcggccgcc |
| | KT164_pMAL_D53M_R | CTTATTTAATTACCTGCAGGGAATTCtcaGATCCTCTGGTGGTCTTGGTG |
| | KT141_D53g-Ex1-2_F | AAGCCAGCTGCAAGCTTGATGGACTCCTTTGTTCCTTT |
| | KT142_D53g-Ex1-2_R | AGGAGTCCATCAAGCTTGCAGCTGGCTTGGAGAAG |
| | KT143_D53g-Ex2-3_F | TTGATCCAGTCAAGGCTAGAGATGATCGGATGGTATT |
| | KT144_D53g-Ex2-3_R | CGATCATCTCTAGCCTTGACTGGATCAAATCCATTGTTA |
| | KT178_D53D1M_325V_F | ACGCACAGCAAGGTCGGCCGCGTCTGGGTGATG |
| | KT179_D53D1M_325V_R | CCAGACGCGGCCGACCTTGCTGTGCGTCTCCAGCA |
| qRT-PCR | OsKUF1_qPCR_F3 | ACTCGGGTTCCTCCTCTGATATAGT |
| | OsKUF1_qPCR_R3 | GAGCCGCTTGTGATTCTGG |
| | qD14L2b-F | CTGCATTGCCTCCATCAACC |
| | qD14L2b-R | AAGCCGCCCTCGTAGTCATC |
| | UBQ RT F1 | AGAAGGAGTCCACCCTCCACC |
| | UBQ RT R1 | GCATCCAGCACAGTAAAACACG |
